# AttSiOff: A self-attention-based approach on siRNA design with inhibition and off-target effect prediction

**DOI:** 10.1101/2023.11.24.568517

**Authors:** Bin Liu, Ye Yuan, Xiaoyong Pan, Hongbin Shen, Cheng Jin

**Affiliations:** Institute of Image Processing and Pattern Recognition, Shanghai Jiao Tong University, Shanghai, China; Medical Robot Research Institute, School of Biomedical Engineering, Shanghai Jiao Tong University, Shanghai, China

**Keywords:** siRNA inhibition prediction, off-target effects, self-attention mechanism

## Abstract

**Motivation:** Small interfering RNA (siRNA) is often used for function study and expression regulation of specific genes, as well as the development of small molecule drugs. Selecting siRNAs with high inhibition and low off-target effects from massive candidates is always a great challenge. Increasing experimentally validated samples prompt the development of machine-learning-based algorithms, including Support Vector Machine (SVM), Convolutional Neural Network (CNN), and Graph Neural Network (GNN). However, these methods still suffer from limited accuracy and poor generalization to design both potent and specific siRNAs.

**Results:** In this study, we propose a novel approach for siRNA inhibition and off-target effect prediction, named AttSiOff. It combines self-attention-based siRNA inhibition predictor with an mRNA searching package and an off-target filter. The predictor gives the inhibition score via analyzing the embedding of siRNA and local mRNA sequences, generated from pre-trained RNA-FM model, as well as other meaningful prior-knowledge-based features. Self-attention mechanism can detect potentially decisive features, which may determine the inhibition of siRNA. It captures global and local dependencies more efficiently than normal convolutions. The 10-fold cross-validation results indicate that our model achieves a significant improvement of correlation between prediction and label, compared with all existing methods. And it reaches better performance of generalization and robustness on cross-dataset validation. In addition, the mRNA searching package could find all mature mRNAs for given gene name from GENOMES database, and the off-target filter can calculate the amount of unwanted off-target binding sites, which affects the specificity of siRNA. Experiments on mature siRNA drugs show that our entire framework, AttSioff, have excellent convenience and operability in practical applications.

## 1 Introduction

RNA interference (RNAi), also known as post-transcriptional gene silencing, can resist to parasitic and pathogenic nucleic acids, and regulate specific gene expression. It have been developed to a mature technology to mediate gene expression manually and probe gene function [1].

RNAi-based regulators include 21∼23-nucleotide small interfering RNA (siRNA) and ∼22-nucleotide microRNA (miRNA). In this paper, we mainly discuss the function and prediction of siRNA. With the help of AGO and TRBP proteins, the antisense strand (AS) in siRNA duplex will bind with target mRNA by Watson-Crick base pairing. If the entire AS can hybridize with target mRNA, it will introduce mRNA cleavage to prohibit the translation process. If only the seed region of AS hybridize with target mRNA, it will induce mRNA degradation and repress the translation process [2] [3] [4] [5].

Usually, target mRNA is composed of hundreds or thousands of nucleotides, from which we can generate a massive amount of siRNA candidates by sliding window method. However, the knockdown efficiency of siRNA, also called inhibition, may vary a lot with a slight change of its composition [6]. The silencing inhibition is mainly determined by the sequence patterns, binding affinity, and the secondary structure around the binding regions, while the specificity is mostly determined by off-target effects [7]. Compared with off-target effects, inhibition is more difficult to predict. All the time, researchers have been focusing on the challenges to predict the inhibition of siRNA accurately.

Early on, methods are mainly developed from rather small datasets, which often contains biased information. For example, Amarzguiouti *et al*. perform a statistical analysis of only 46 siRNAs to identifying the preferences of nucleotides on each position of siRNA duplex. They find that the motifs U1 or G19 is strongly related to poor inhibition [8].

With the growth of valid samples, algorithms based on machine learning have been developed in a data-driven way. Heusken *et al*. make the greatest contribution to enrich the relevant dataset. They collect 2431 siRNAs targeting 34 mRNAs with corresponding validated inhibitions. And they develop a model named BIOPREDsi using Stuttgart Neural Net Simulator to achieve a rather high Pearson Correlation Coefficient (PCC) of 0.66 [9]. Vert *et al*. perform a LASSO-based regression model, to conveniently estimate the importance of each feature. They use the preference for specific nucleotide on some positions, as well as short asymmetric base motifs, as the input [10]. Ichihara *et al*. develop a simple linear regression algorithm, i-Score, which is only comprised of nucleotide preferences at each position as the input, and achieves a comparative PCC with s-Biopredsi [11]. However, these methods still suffer from incompetence of detecting hidden features.

In recent years, Convolutional Neural Network (CNN) has been successfully applied to diverse fields, such as machine translation, object detection, protein interaction, etc. It has also been used in siRNA inhibition prediction and showed remarkable enhancement of precision. Similar to TextCNN, the work done by Han *et al*. utilizes multiple convolutional kernels to detect unknown but helpful motifs from local target mRNA sequence, preprocessed by one-hot encoding. Then it uses average pooling and maximum pooling layers to extract the most representative features. The thermodynamic property calculated from AS is concatenated with the pooling output, and then normalized in batch. At last, a neural network with one hidden layer of 25 nodes is applied, and this model generates the prediction score via sigmoid activation function [12]. It achieves a remarkable improvement on PCC compared with traditional models. However, the inadequate input features and simple hidden layer limit its performance. And the forceful pooling operations result in great loss of information.

Aside from CNN, Graph Neural Network (GNN) is another common deep learning algorithm used in bioinformatics. Biological molecules are regarded as nodes, and their relationships can be represented with edges connecting different nodes [13]. Graph is an intrinsically good structure to model topology and capture hidden interrelationship in non-structural data. Massimo *et al*. propose a GNN-based model for siRNA inhibition prediction for the first time. There are three types of nodes in their graph. The first is siRNA node, with 3-mer counting as its feature. The second is target mRNA node, with 4-mer counting as its feature. And the third is siRNA-mRNA interaction node, with the thermodynamic parameters calculated from Gibbs energy and RNAup program as its features. Here, they replace the interaction edge with node, for the sake of thoroughly incorporating the interaction information into the graph. And they consider the inhibition as the property of siRNA-mRNA interaction node to predict. This algorithm reaches a better performance than aforementioned CNN-based method. However, the k-mer counting features are still insufficient to represent the characteristics of siRNA or mRNA sequence. And the graph is fixed with nodes from trainset and test set, resulting in its disability to predict the inhibition of new siRNA samples [14].

Despite that various researches have been developed to predict the siRNA inhibition, there is still room for developing a new algorithm with better accuracy and generalization. This can be achieved by optimizing the feature selection and modeling process. Besides, though many studies in vitro have shown the capacity of siRNAs in gene silencing, some challenges still exist before applying siRNA into clinical trials, such as limited longevity and inevitable off-target effects[15] [16] [17]. Off-target effects will result in serious misjudgment of inhibition. And silencing uncertain mRNAs may negatively interfere with some significant biochemical pathways. Compared with difficult inhibition prediction, off-target effect is easier to analyze with some definite criteria.

In this study, we propose a novel approach for siRNA inhibition and off-target effect prediction, named AttSiOff. This self-attention-based inhibition predictor employs two types of features. One is the embedding of siRNA and local target mRNA sequences, generated from pre-trained RNAFM model. The other is prior-knowledge-based characteristics of AS, including the thermodynamic parameters, the secondary structure, GC content, PSSM score, etc. And this predictor is comprised of two parts: feature extraction module and fully connected module. In the feature extraction module, multi-head self-attention mechanism is used to further extract hidden information by constructing the interaction of every nucleotide with others. To the best of our knowledge, it is the first time of self-attention mechanism to be used in the prediction of siRNA inhibition. In the fully connected module, the high-dimensional representations produced by multi-head self-attention are concatenated with other features and go through a deep and wide neural network, to produce a prediction score. The 10-fold cross-validation results and cross-dataset experiments both show that our predictor achieves the state-of-the-art performance, compared with other existing methods.

To facilitate and streamline siRNA design, we combine the predictor with an mRNA searching package and an off-target filter. The mRNA searching package can find all mature mRNAs for any given gene name. The off-target filter can calculate the amount of possible unwanted off-target binding sites, which affects the specificity of siRNA. We testify the practicability and maneuverability of our pipeline on five siRNA drugs from Alnylam Company. The results show its great simplicity and user-friendliness, and verify the utility of our inhibition predictor too.

We proclaim the following contributions in our approach: (1) We apply the pre-trained RNA-FM model to greatly enrich the embedding of the RNA sequences, instead of using classic one-hot binary encoding method. (2) We successfully employ self-attention mechanism in the work of siRNA inhibition prediction for the first time to capture the global and local dependencies. (3) Our predictor achieves the best performance on both prediction accuracy and cross-dataset generalization, compared with other methods. (4) We construct a simple and user-friendly approach to automatically design both potent and specific siRNAs.

## 2 Materials and Methods

### 2.1 Related work

Attention mechanism can efficiently capture the long-distance dependencies, compared with CNN or Recurrent Neural Network (RNN). It is first applied in computer vision, and combined with RNN to classify images [18]. Then it is employed in machine translation to form an en-coder-decoder architecture. The encoder encodes a source sentence into a pre-defined fixed-length vector, and the decoder will generate a target sentence accordingly. Attention mechanism helps a lot to solve the problem of limited receptive field or difficult alignment in classic CNN or RNN [19]. The most significant work done by Vaswani *et al*. discards the recurrence and convolutions totally, and only utilizes multi-head self-attention mechanism to capture interaction relationship. The well-known TransfomerEncoderLayer contains two sublayers, and both of them are formed in residual connection structure. The first sublayer is a multi-head self-attention mechanism. And the second sublayer is a simple fully connected feed-forward module [20].

Understanding the structure and function of RNA sequence is essential to help explain some biological pathways, such as post-transcriptional regulation, and cell signaling. With the increasing development of sequencing technology, more and more RNA sequences are probed. However, annotated RNA sequences are very limited. To address this problem, Chen *et al*. propose a transformer-based model, named RNA Foundation Model (RNA-FM), to train on 23 million ncRNA sequences from RNAcentral database via self-supervised learning. The clustering results indicate that the pre-trained RNA-FM embedding contains sequential and functional information. And it achieves better performance when applied to downstream tasks, such as secondary structure prediction, and 3D affinity prediction [21].

### 2.2 Datasets

As is suggested in the work of Han *et al*. [12], we try to obtain as many experimentally validated siRNAs as possible. In this study, 3536 siRNAs from the work done by Huesken [9], Reynolds [22], Vickers [23], Haborth [24], Takayuki [25], and Ui-Tei [26], are collected. We divide these samples into three datasets according to their experimental conditions, namely DH, DR, and DT. The detailed composition of these three datasets is shown as **Table 1**.

**Table 1.**
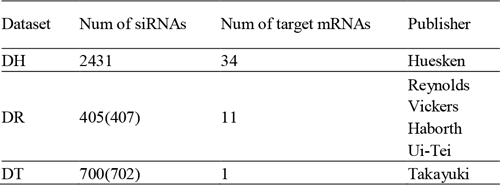
The details of three datasets. Two siRNAs are removed in DT due to the limitation of i-score website, and two are discarded in DR as a result of lacking target binding sites.

Two siRNAs in DR are removed due to failure to find target binding sites. And two siRNAs in DT are discarded due to the limitation of i-Score website, which will be explained later. As is said above, the experimental conditions of these datasets are very different. Their inhibition labels range from 0 to 134.1, -27.8 to 98.9, and 0 to 97, respectively. To unify the data distribution, we normalize their inhibition labels individually before combining them together as a whole dataset, named DHRT. And we sort the samples in DHRT in an inhibition-decreasing order.

Generally, only the core region (19 nucleotides from the 5’ end) of AS will hybridize with target mRNA. After collecting those siRNA samples, we refer to the publications to get corresponding local target mRNA sequences, which are defined as 19 nucleotides on the binding site plus 20 downstream and 20 upstream nucleotides along the target mRNA (59 nucleotides totally). This number of nucleotides in flanking regions comes from the idea of Han *et al*. [12].

In addition to the published datasets, we collect five siRNA medicines from Alnylam Company, which are applied to clinical and diagnostic usage in recent years (**Supplementary Table S1**). Although these siRNAs are chemically modified to strengthen the potency, prolong the longevity, and weaken off-target effects, we can remove the chemical components here and use the bare sequences to further validate the robustness and generalization of our inhibition predictor, as well as the practicability and maneuverability of our approach.

### 2.3 The architecture of our predictor

As is shown in **Fig. 1**, our predictor consists of feature extraction module and fully connected module. And two types of features are employed as input. One is the encoding of siRNA and local target mRNA sequences, generated from pre-trained RNA-FM model. The other is prior-knowledge-based characteristics of AS, including thermodynamic parameters, k-mer counting, PSSM score, the secondary structure, and GC content.

**Fig. 1:**
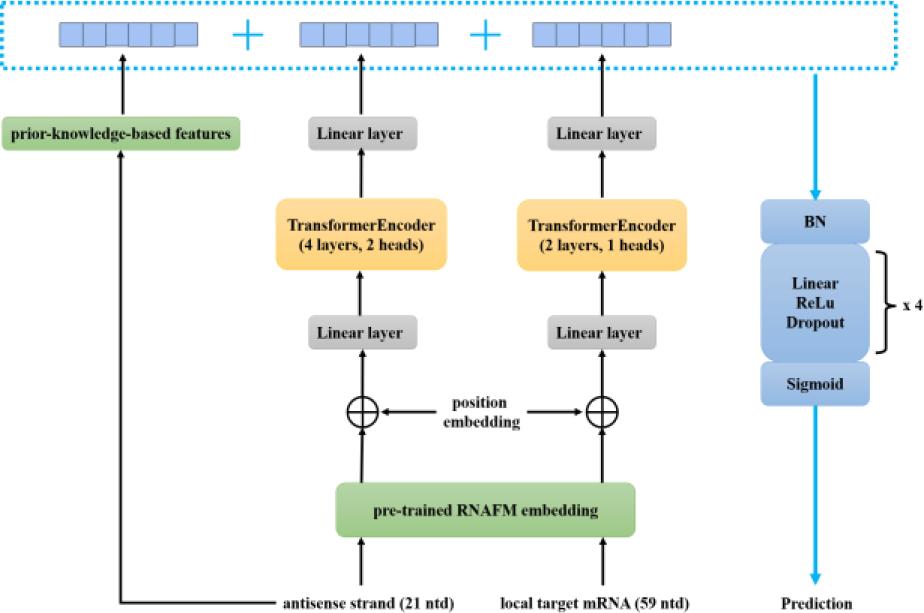
The architecture of our self-attention-based siRNA inhibition predictor. Two kinds of features are utilized. One is the pre-trained RNAFM embedding. The other is prior-knowledge-based features. The TransformerEncoder extracts hidden features from the RNAFM embedding, and the fully connected module completes the feature fusion and gives the prediction.

#### 1) Encoding of sequence context

In this study, we consider the embedding of pre-trained RNA-FM model as the representation of siRNA and local mRNA sequence context, instead of using simple one-hot encoding method. RNA-FM model will transform each nucleotide into a 640-dimensional continuous vector, which proves to contain structure-related and function-related information. We believe it will enrich the sequence representations greatly. In contrast, one-hot encoding method will transform each nucleotide into a 4-dimensional binary vector here. It is known for its disability in modeling the correlation relationship between different kinds of nucleotides. Besides, it ignores the dependencies among nucleotides in adjacent positions.

Note that not all local target mRNA has complete flanking region around its binding site, so we use a special vector [0.05] * 640 to represent the missing nucleotide in RNAFM embedding.

#### 2) Other prior-knowledge-based characteristics

Although the following prior-knowledge-based characteristics are mainly described in previous researches from biased dataset, they do play a role in predicting the inhibition of siRNA. And our fully connected module can automatically learn to distinguish those useless features from decisive ones. In this paper, we mainly consider five groups of relevant features.

The first one is thermodynamic stability. Relative study has shown that the stability of siRNA duplex has an importance on its inhibition and longevity, as well as determining which strand in siRNA duplex will hybridize with target mRNA [25] [2] [27] [28]. We use the Gibbs free energy to describe the thermodynamic characteristics of the core region of AS **(Supplementary Table S2)**. There are 20 parameters in total, including 18 features calculated from every two adjacent bases, 1 feature from the difference between its 5’ end and 3’ end, and 1 feature of overall energy.

The second one is k-mer counting. We compute the frequencies of 1-mer, 2-mer, and 3-mer segments to represent the presences of 4, 16, and 64 kinds of possible short motifs.

The third one is PSSM score. Position Specific Scoring Matrix (PSSM) describes the possibility of observing one nucleotide on each position. Here, we generate the PSSM from statistical analysis of the entire train-set. And for any new siRNA in test set, we utilize the fixed PSSM to estimate the similarity between the new siRNA and all samples in train-set.

The forth one is related to secondary structure. Previous work has indicated that unstructured AS can mediate more active gene silencing. That means folded antisense strand is hard to hybridize with target mRNA, consequently reducing its inhibition [29]. Here we use RNAfold program to predict the minimum free energy and possible base pairing percentage as the secondary-structure-related features [30].

The last one is GC content. They mainly affect the inhibition by changing the thermodynamic property of siRNA, because the stability between base G and C is much stronger than that between base A and U. The higher the GC content is, the more stable siRNA is [31]

#### 3) Feature extraction module

The feature extraction module is composed of one linear layer, several continuous TransformerEncoderLayer (4 layers for siRNA, and 2 layers for local target mRNA), and another linear layer. Normal convolution is hard to capture the long-distance dependencies, unless the convolutional kernel is big enough to cover the whole sequence. Thus, we consider self-attention mechanism to construct the interaction relationship among all nucleotides, to automatically capture hidden decisive motifs. However, the initial dimension of embedding processed by RNA-FM is too big to extract valid information. It may contain many redundant and task-irrelevant features, and the high-dimensional input will overburden the following self-attention mechanism. Therefore, the first linear layer is used to squeeze it to form an 8-dimensional high-level representation, and reduce the complexity and computation of subsequent layers. The RNN or CNN usually can capture the relative and obsolete position information via striding ahead along time or feature matrix, while attention mechanism has no ability to distinguish the sequential order. We add the sine and cosine position embedding into the embedding, as indicated in the work of Vaswani [20]. Then following TransformerEncoderLayer will capture the interconnected hidden motifs easily. The second linear layer at the end of feature extraction module play the similar role of pooling. It will take all features into account, mapping the 8-dimensional vector into 1-dimensional vector, instead of just selecting the maximum or average value.

#### 4) Fully connected module

The fully connected module is composed of a batch normalization layer, four non-linear sublayers, and a sigmoid activation layer. Each non-linear sublayer consists of a linear layer, a ReLU activation function, and a dropout layer. Input features are made up of the output of feature extraction module, and other prior-knowledge-based features. We normalize the concatenation in batch first, for the sake of faster convergence and better generalization. The overall feature vector is then fed into the four sublayers to complete the feature fusion. The sigmoid activation function is used to generate the prediction score of siRNA inhibition.

### 2.4 The architecture of our approach

To facilitate and streamline siRNA design, we construct an approach, named AttSiOff. Aside from the aforementioned siRNA inhibition predictor, it consist of an mRNA searching package and an off-target filter.

#### 1) The mRNA searching package

Usually, probed mRNA sequences may update with the development of molecular biology, and only target gene name is provided in siRNA design. Downloading mRNA sequences manually on NCBI or other websites is time-cost and annoying. Fortunately, we find one package, pyGB, implemented by Haotian Teng. For any given gene name, it could search corresponding mature mRNAs rapidly.

#### 2) The off-target filter

To build an effective siRNA design tool, we shall consider the off-target effects, which may weaken the siRNA inhibition, intervene normal necessary cell activities, and do harm to receptors. They may arise from three aspects: on-target silencing of unintended mRNAs (perfect or near perfect base pairing with untargeted mRNAs), miRNA-like off-target silencing (imperfect base pairing with 3’ UTRs of unwanted mRNAs), and stimulation of innate immune response [32] [33].

For on-target unintended silencing, it is caused by 16 or more consecutive base pairings between the AS and unwanted mRNAs. The 19-nucleotide SS can be segmented into four 16-nucleotide subsequences, and then be searched using substring searching algorithm. However, some effective sites do not confirm to perfect canonical seed pairing, which may allow for wobble or mismatch [34]. To solve this problem, we can use Smith-Waterman algorithm, which calculate optimal alignment between two sequences. One siRNA may have multiple binding sites on one mRNA sequence. Thus we need to make some improvements for detecting multiple outputs, by replacing the one-off backtracking with recurrent backtracking, until the current maximum score is less than a specific threshold.

MicroRNA-like off-target effect refers to siRNA-induced regulation of unintended transcripts, through partial sequence complementarity to their 3’UTRs [32]. The most possible binding sites for miRNA are: 8mer site (base pairing at positions 2-8 with a base A opposite at position 1), 7mer-m8 site (base pairing at positions 2-8), and 7mer-A1 site (base pairing at positions 2-7 with a base A opposite position 1) (**Supplementary Fig. S1**) [34]. We can also search these three kinds of target sites by substring searching.

As for the non-specific immune response caused by siRNA, it can be reduced by selecting bases carefully to avoid containing putative immunostimulatory motifs UGUGU and GUCCUUCAA in the AS [35]. Thus, we check if each siRNA contains the two motifs by substring searching.

#### 3) The flow of our approach

The architecture of our siRNA design approach is shown as **Fig. 2**. The entire workflow is divided into four parts.

**Fig. 2:**
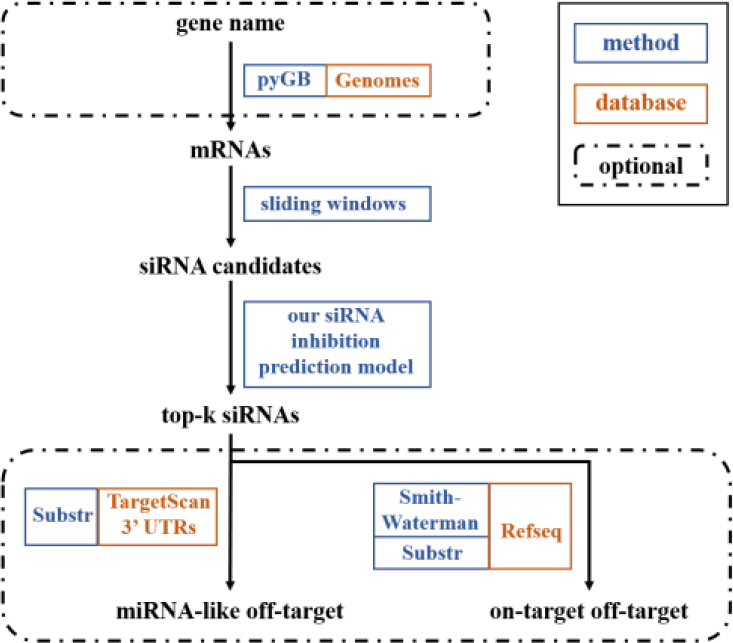
the architecture of our siRNA design approach, HySioff. It contains the mRNA searching package pyGB, siRNA inhibition prediction model, and off-target filter.

First, the pyGB package is used to search for mature mRNAs according to the input gene names from GENOMES database. One can also provide mRNA sequences in FASTA format directly.

Second, every mRNA sequence is cleaved to generate all alternative 19-nucleotide SSs via sliding window, and get corresponding 21-nucleotide ASs, 59-nucleotide local target mRNA, and other attributes simultaneously. Considering the potential weak specificity and possible toxicity, the following characteristics in the AS are also collected in this step: the Gibbs free energy, presence of two immunostimulatory motifs, presence of long stretches of identical bases, GC content, GC content, presence of more than 2 continuous CAN pattern, and presence of more than 2 consecutive CUG/CCG/CGG motifs [36] [37].

Third, every siRNA duplex and opposite local mRNAs are fed into our siRNA inhibition predictor, and all siRNAs targeting identical gene are sorted in prediction-descending order.

Forth, one can choose optional off-target filter for top-k siRNAs. Here, we mainly take use of two databases: Refseq (downloaded from ftp://ftp.ncbi.nih.gov/refseq/H_sapiens/mRNA_Prot/human.rna.gbff.gz) and TargetScan (downloaded from https://www.targetscan.org/cgi-bin/targetscan/data_download.vert72.cgi). As is discussed above, the miRNA-like off-target and on-target off-target can both be calculated using substring searching. And the on-target off-target can also be predicted by improved Smith-Waterman algorithm.

This workflow will eventually provide the user with top-k siRNAs with their multiple attributes and possible off-target effects, which facilitate the pre-screening stage greatly.

### 2.5 Experimental Setup

To ensure the uniform distribution between trainset and test set, we sample the siRNA with indices of i, i+10, i+20…in DHRT to form the test set during 10-fold cross-validation. As for cross-dataset validation, we use different sources of datasets as the trainset and test set.

Our method is built with Pytorch in python. We choose the Adam optimizer with weight decay of 5e-4 and initial learning rate of 0.01. Drop-out operation exists both in the multi-head self-attention module and the fully connected module, to repress the impact of overfitting. In addition, we set the maximum epoch to 1000. And we use early-stopping strategy to supervise the PCC metric. If this indicator continues decreasing for 20 epochs, it will terminate the training phase, and the model parameters with the best PCC will be saved. As a regression task, we train the model with the mean square error (MSE) loss.

### 2.6 Evaluation metrics

To estimate the prediction performance, we use three statistical indicators here, including Pearson correlation coefficient (PCC), Spearman correlation coefficient (SPCC), and the Area under the Receiver Operating Characteristic curve (AUC). PCC and SPCC evaluate the linear correlation between two sets of data. AUC is used to evaluate the performance of binary classification. In this paper, we use 0.7 as the inhibition threshold to classify a siRNA to be positive or negative.

Among these three metrics, SPCC is the most important one. It denotes the correlation of rankings between predictions and labels. In siRNA design, precise ranking will reduce the workload to find functional siRNAs.

## 3 Results and discussion

### 3.1 10-fold Cross-validation result

We compare our model with i-Score [25], Biopredsi [9], DSIR [10], one CNN-based model [38], and one GNN-based model [39]. The first three algorithms lack of source code, and they are hard to reproduce. Fortunately, we find the i-Score webserver can generate all siRNA candidates with predictions of these models, for any given target mRNA sequence. But siRNAs at the first two positions are limited to predict, and that is why we discard two samples in DT above.

The 10-fold cross-validation result is shown as **Fig. 3**. Apparently, our model achieves the state-of-the-art performance among the six methods, reaching an average PCC of 0.7612, SPCC of 0.786, and AUC of 0.898.

**Fig. 3:**
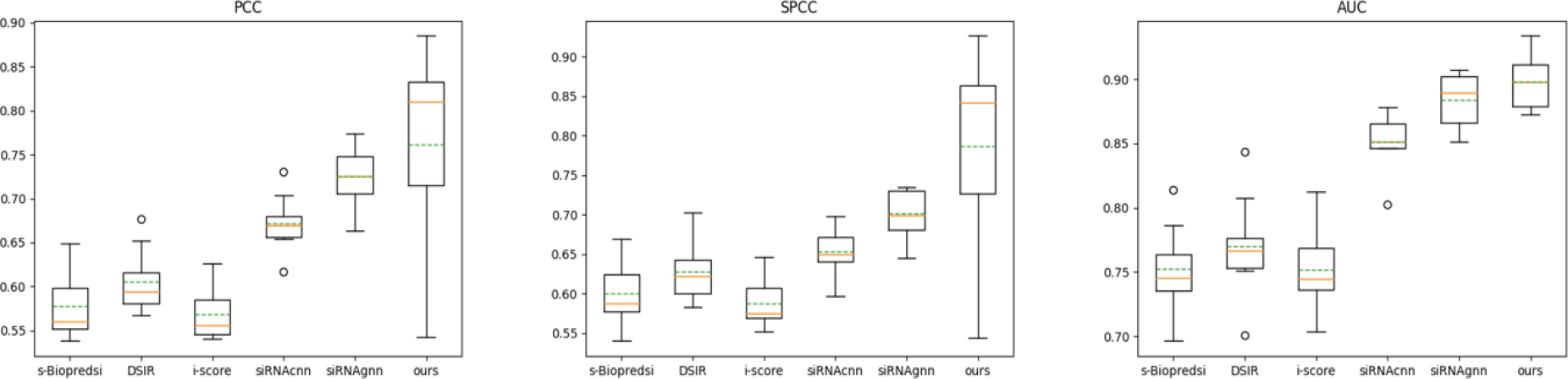
The 10-fold cross-validation result of our predictor compared with existing methods. Three metrics are PCC, SPCC, and AUC, respectively.

In comparison, the three traditional methods show poor performance on all indicators. The reasons may be that their inputs, based on manual feature engineering, lack significant information and are usually biased. And their models have limited predicative capability to capture hidden motifs.

The CNN-based model reaches an average PCC of 0.669, SPCC of 0.647, and AUC of 0.853, which are much lower than ours. We may deduce that their convolutions with multiple kernels only focus on local adjacent correlation, and fail to capture global interrelationship of the entire sequences. And the forceful pooling operation results in loss of significant information. Most importantly, their one-hot encoding representations suffer from the aforementioned problems, the sequence features obtained from which contain scanty information.

The performance of GNN-based model is not bad, achieving an average PCC of 0.725, SPCC of 0.7, and AUC of 0.884. The idea of modelling siRNA-mRNA interaction task with graph structure is innovative and intuitive. Their experiments demonstrate that the biological graph model have potent predictive capability, even with limited features as the property of nodes. Their predictive accuracy may further improve with more meaningful features as input. However, the most serious problem of the GNN-based model is that it cannot predict on new siRNA sample with a fixed graph. That is why we exclude this model in the following cross-dataset validation.

### 3.2 Cross-dataset validation

For better demonstration of generalization of our method, we evaluate the cross-dataset performance. We train the model only on DH, DR, and test on DT, or train it only on DH, DT, and test on DR. These three datasets are totally independent from each other. The results are shown in **Table 2**. It can be expected to notice a big drop in all metrics compared with those of 10-fold cross validation, due to out-of-distribution problem.

**Table 2.**
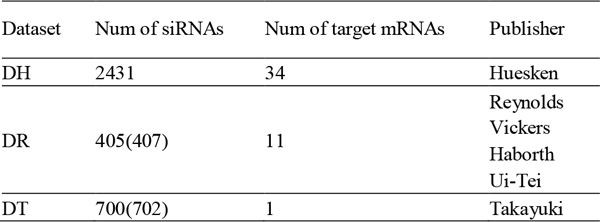
The cross-dataset prediction results. Bold numbers indicate the best results.

Although the number of parameters in our model is much bigger than existing methods, it does not show the problem of overfitting. And it still significantly outperforms all other methods on cross-dataset validation, beyond our expectation. We may deduce that the batch normalization, dropout, and early-stopping strategy all help to ensure a good generalization.

To interpret the internal reasons why all indicators on DT are higher than those on DR, we use T-distributed Stochastic Neighbor Embedding (t-SNE) to visualize the RNA-FM embedding of siRNA sequences from three datasets **(Supplementary Fig. S2)**. The samples in DT are intrinsically divided into three clusters, while the samples in DR distribute more chaotically. That means predicting on DT is easier than that on DR coherently. We also analyze the differences of the inhibition distribution between these three datasets (**Supplementary Fig. S3**). The kernel density estimate (KDE) plots show that the distributions of DH and DT are quite similar, but they differ from DR. This may be another reason why all models perform better on DT than DR

### 3.3 Comparison on five siRNA drugs

Moreover, we evaluate our method on the aforementioned five siRNA drugs. The reported median knockdowns, the predicted inhibitions, the rankings of predictions among all corresponding siRNA candidates, and the number of possible off-target binding sites are shown in **Supplementary Table S3**. Theoretically, the inhibition of bare sequence is supposed to be less than that of chemically modified siRNA.

Our approach, AttSiOff, helps a lot to facilitate this experiment. First, we take the target gene names as the input, namely TTR, ALAS1, PCSK9, and HAO1. The mRNA searching package then automatically searches latest mature mRNA sequences from GENOMES database and collects all possible siRNA candidates by sliding window. We do not need to download those mRNA sequences manually. Second, our predictor predicts the inhibitions of siRNAs grouped by gene names, and sorts them in inhibition-descending order. We use the location of these five siRNAs in their respective candidate sets as the ranking score. Third, for the five siRNAs, the off-target filter will compute the number of possible off-target binding sites.

The rankings show that inhibitions of the five siRNAs, predicted by our model, rank near the top in all candidates (1.79%, 0.62%, 2.4%, 11.2%, and 3.58%), which means researchers need fewer experiments to find wanted drugs, compared with other methods. And the predicted off-target effects are in allowable range, to ensure the specificity.

The comparison results demonstrate that our predictor outperforms other methods and can facilitate the design process for given gene.

### 3.4 Ablation study

More experiments are executed on the effect of different types of TransformerEncoder. 2, 4, 8 layers, and 1, 2, 4 heads are tested, separately. The results show that inadequate or excessive layers or heads both decrease the prediction performance. To obtain the optimal PCC metric, we select 4 layers with 4 heads for siRNA embedding, and 2 layers with 1 head for mRNA embedding.

And we test the effect of different loss function. In regression task, MAE and MSE are frequently used. The observed PCC are 0.712 for MAE, and 0.761 for MSE. The reason may be that MSE loss will pay more attention to those anomalous samples to get more stable closed-loop solutions for weights. And this will lead to better generalization on test set.

In addition, we emphasize the advantages of pre-trained RNAFM embedding, compared with one-hot encoding. Thus, we also execute comparative experiments by replacing the RNAFM embedding with one-hot encoding on sequence context. The result gives a 21% average decrease on PCC.

## 4 Conclusions

Hundreds or thousands of siRNAs may target the same mRNA sequence. Utilizing computational methods to identify those hyperfunctional siRNAs from massive candidates has increasingly become a significant study. However, existing methods are still not accurate and robust enough to design potent and specific siRNAs, and most of them have not consider unintended off-target effects.

In this paper, we propose a novel self-attention-based approach on siRNA inhibition and off-target effect prediction, named AttSiOff. First, the mRNA searching package helps obtain target mRNA sequences for given gene name. Then the inhibition-related hidden features are captured from pre-trained RNA-FM embedding through multi-head self-attention mechanism. After being concatenated with other prior-knowledge-based features, they are fed into the fully connected module to complete the feature fusion and give the inhibition prediction. At last, the off-target filter utilizes substring searching or improve Smith-Waterman algorithm to give the possible amount of off-target binding sites. Compared with existing methods, our approach shows four major advantages. First, we include prior-knowledge-based features as input, such as GC content, the secondary structure, etc. These features analyzed from small biased dataset still contribute to precise prediction. Second, we use pre-trained RNA-FM model to encode the sequence context. The high-dimensional embedding contains more meaningful information than one-hot binary encoding, especially the functional and structural information. Third, we replace convolution with multi-head self-attention mechanism, to capture the global long-distance dependencies within the entire sequence. Forth, we use additional filter to predict the off-target effects, to further ensure its specificity in practical application.

To evaluate the validity of our predictor, relevant comparison experiments are designed for verification. Experimental results show that our model achieves state-of-the-art performance on PCC, SPCC, and AUC, compared with classical methods based on ANN, LASSO, SVM, CNN, and GNN. And the cross-dataset validation demonstrates the brilliant generalization and robustness of our model. Besides, our automatic siRNA design tool, AttSiOff, facilitates our experiments on five siRNA drugs, and we hope it can help other researchers, who are devoted to designing both effective and specific siRNA.

This study provides new perspectives and analytical ideas for siRNA inhibition and off-target effects prediction. And we hope it help bring attention mechanism to broader bioinformatics-related applications. Since some chemically modified siRNAs are deliberately designed to further improve their inhibitions and reduce the off-target effect, AttSiOff will be used as backbone and the representations of chemical modification will be taken into account in the future.

## Abbreviations

RNA-FM: RNA fundamental model
RNAi: RNA interfering
siRNA: small interfering RNA
CNN: Convolutional Neural Network
GNN: Graph Neural Network
PCC: Pearson Correlation Coefficient
SPCC: Spearman Correlation Coefficient
AUC: the Area under the Receiver Operating Characteristic curve

## Declarations

### Availability of data and material

Not applicable.

### Authors’ contributions

Conceptualization and original draft preparation, Bin Liu; reviewing and editing, Bin Liu, Ye Yuan; Ye Yuan, Xiaoyong Pan, Hongbin Shen, and Cheng Jin revised and provided critical feedback of the manuscript. All authors have read and agreed to the final version of the manuscript.

## Acknowledgements

Not applicable.

